# Transcriptional activation of Il36A by C/EBPβ via a half-CRE•C/EBP element in murine macrophages is independent of its CpG methylation level

**DOI:** 10.1101/2021.04.24.441265

**Authors:** Andreas Nerlich, Nina Janze, Ralph Goethe

## Abstract

Interleukin-36α(Il-36α) is a member of the novel Il-1-like proinflammatory cytokine family that is highly expressed in epithelial tissues and several myeloid-derived cell types. We have recently shown that CCAAT enhancer binding proteinβ (C/EBPβ) binds specifically to an essential half cAMP response element (half-CRE)•C/EBP motif in the *Il36A* promoter to induce *Il36A* expression upon LPS stimulation. C/EBPs are transcription factors belonging to the basic leucine zipper (bZIP) family of transcriptional regulators. C/EBP proteins can form homo- and heterodimers and regulate gene expression by binding to C/EBP specific recognition sequences and composite sites that can contain 5′-cytosine-phosphate-guanine-3′dinucleotides (CpG). CpG methylation of such elements has been shown to influence transcription factor binding and gene expression. Here we show that the half-CRE•C/EBP element in the *Il36A* promoter is differentially methylated in the murine RAW264.7 macrophage cell line and in primary murine macrophages. By using electrophoretic mobility gel shift and fluorescence polarization assays we demonstrate that C/EBPβ binding to the half-CRE•C/EBP element in the *Il36A* promoter following LPS stimulation is insensitive to CpG methylation. Transfection assays also show that methylation of the CpG in the half-CRE•C/EBP element does not alter LPS-induced *Il36A* promoter activity. A direct comparison of *Il36A* mRNA copy numbers as well as the pro-Il-36α protein level in RAW264.7 and primary macrophages revealed similar amounts in both cell types. Taken together, our data suggest that C/EBPβ binding to the half-CRE•C/EBP element and C/EBPβ mediated gene activation occurs independently of the CpG methylation status of the target DNA sequence and underline the potential of C/EBPβ to recognize methylated as well as unmethylated binding sites.

## Introduction

The interleukin (Il)-36 cytokines constitute a subfamily of the Il-1 cytokine family and consist of three agonistic cytokines (Il-36α, Il-36β, Il-36γ) and the Il-36 receptor antagonist (Il-36RA) that play important roles in host immunity (1). Il-36 cytokines are expressed by a variety of cells in different tissues, such as macrophages, dendritic cells, keratinocytes, and lung epithelial cells (2–7). They bind to the Il-36 receptor (also referred to as Il-1R-like 2) that is widely expressed on many different cells, including murine bone marrow-derived dendritic cells, CD4^+^T cells, mononuclear phagocytes, and various epithelial cells of skin, lung, and digestive track tissues (2,8,9). Similar to the receptor for Il-1, the Il-36 receptor recruits the transmembrane Il-1 receptor accessory protein upon ligand binding that subsequently activates intracellular signaling via mitogen-activated protein kinases and nuclear factor-ϰB (NF-ϰB) activated transcription leading to inflammatory responses (10,11). At the transcriptional level Il-36α and Il-36γ themselves are regulated by NF-ϰB, C/EBPβ and by the transcription factor ‘T-box expressed in T cells’ (T-bet), respectively (3,5).

DNA methylation at cytosine residues by DNA methyltransferases is an epigenetic modification involved in mammalian development, lineage identity as well as transcriptional regulation (12). Traditionally, DNA methylation of gene promoters was considered to silence transcription either directly by inhibiting binding of certain transcription factors (TF) to their binding sites (13–16) or indirectly by binding of TFs containing a methyl-CpG (mCpG)-binding domain (MBD), that in turn recruits histone deacetylases which subsequently promote local chromatin condensation (17). In recent years, a growing number of TFs lacking MBDs have been shown to bind to their transcription factor binding sites (TFBS) even when they are methylated, arguing against the paradigm that CpG methylation always represses transcription. Examples include transcriptional activators that contain a C2H4 zinc-finger domain like Kaiso and Krueppel-like factor 4 (18,19), proteins with a helix-turn-helix DNA-binding domain like Zhx1/2 and Pax6 (20,21), helix-loop-helix domain containing TFs like c-MYC and ARNT2 (22,23), and TFs belonging to the family of bZIP proteins like C/EBPs (24,25).

C/EBP transcription factors belong to the bZIP family of transcription factors and are involved in tissue-specific gene expression, proliferation, differentiation, and inflammation (26). In particular, C/EBPβ has been shown to regulate inflammatory genes like the cytokines Il-6, Il-12 p40, and Il-36α (5,27,28), the chemokines Il-8 and macrophage inflammatory protein-1α (27,29) as well as the proinflammatory genes for inducible NO synthase (NOS2) and cyclooxygenase-2 (30,31) in macrophages. DNA binding and subsequent gene expression by C/EBPβ requires formation of homodimers or heterodimers with other C/EBP family members or members of the CREB/ATF family via the bZIP domain (32). The consensus sequence as well as composite sites recognized by C/EBPs can contain a central CpG dinucleotide (26). Several studies showed that C/EBPs are able to bind these sequences *in vitro* and *in vivo* with similar or even with increased affinities if the central CpG dinucleotide is methylated (16,24,25,33). As a consequence, such methylation can generate C/EBP binding sites at CRE-like sequences leading to activation of a subset of differentiation specific genes by C/EBPs whilst inhibiting activation of these genes by CREB members (24). The structural basis for this insensitivity to the methylation status of the bound DNA has been recently unravelled and was shown to be mediated via a so-called methyl-Arg-G triad (34).

Recently, we reported that C/EBPβ confers transcriptional activation of *Il36A* via an essential half-CRE•C/EBP site that contains a central CpG dinucleotide (5). For Il-36 cytokines, there is to our knowledge, no information on epigenetic regulation, in particular on CpG methylation and its possible impact on transcription factor binding, available. Therefore, we herein analyzed the methylation level of this element in two different murine macrophage types and its impact on C/EBPβ binding and transcriptional activation of *Il36A*.

## Materials and Methods

### Reagents

Media used for macrophage cell culture were obtained from ThermoFisher Scientific (Darmstadt, Germany) and X-tremeGENE reagent was obtained from Roche (Mannheim, Germany). Purified ultra-pure LPS from *Escherichia coli* 0111:B4 and chemically competent *E. coli* GT115 were purchased from Invivogen (Toulouse, France). If not stated otherwise all other reagents were from Sigma (Taufkirchen, Germany). Antibodies against ATF/CREB-1 (Cat# sc-270, RRID:AB_2290030), CREB-2/ATF-4 (Cat# sc-200, RRID:AB_2058752), C/EBPβ (Cat# sc-150, RRID:AB_2260363), and C/EBPδ (Ca# sc-151, RRID:AB_2078200) were purchased from St. Cruz (Heidelberg, Germany). The antibody against GAPDH (D16H11, RRID:AB_11129865) was obtained from Cell Signaling (Frankfurt am Main, Germany) and goat anti-mouse Il-36α (AF2297, RRID:AB_355216) was from R&D Systems/Bio-Techne (Wiesbaden-Nordenstadt, Germany). Secondary goat anti-rabbit IgG, HRP-linked (#7074, RRID:AB_2099233) was obtained from Cell Signaling and donkey anti-goat IgG AP-linked (Cat# sc-2022, RRID:AB_631723) was from St. Cruz.

### Plasmid construction and *in vitro* methylation of plasmid DNA

The Plasmid pCpGL-TA-Il36A-357/-45 was generated by cloning a synthetic DNA fragment containing the TATA box of the human EF1 promoter (5′-GATCTCAG GGTGGGGGAGAACCATATATAAGTGCAGTAGTCTCTGTGAACATTCA-3′) into pCpGL-basic (kindly provided by M. Rehli). Subsequently the *Il36A* promoter region from postiion −357 to −45 relative to the transcriptional start site (TSS) was amplified using the oligonucleotides PromIL36A_for-357 (5′-GCCTGCAGTTGCACTTCCTGTAGGTTC-3′) and PromIL36A_rev-45 (5′-GCA GATCTAGAGGAGGTTATGCCTCAG-3′) and the plasmid pGL3-Il36A-1120-Luc (5) as template. All pCpGL plasmids were maintained in *E. coli* GT115. Plasmid pJet-Il36A was generated by amplifying a fragment of *Il36A*from cDNA using the qRT-PCR primers Il36A_for (5′-AAGGAACCTGTAAAAGCCTCTCT-3′) and Il36A_rev (5′-CAGTTCTTGGGTCAGAATGAGTG-3′) and subsequent cloning into pJET1.2/blunt vector as described by the manufacturer (ThermoFisher Scientific). The DNA fragment encoding mouse C/EBPβ residues 221–296 (bZIP domain) was cloned into pET-45b(+) vector (Novagen/Merck, Darmstadt, Germay) via ‘*in vivo* assembly’ (35) to generate a His-tagged version of the bZIP domain. The bZIP domain was amplified using the primers pET-45b_f (5′-CTGGTAAAGAAACCGCTGCTG-3′) and pET-45b_r (5′-GTGATGGTGGTGGTGATGTG-3’) and the plasmid pcDNA-C/EBPβ-wt (5) as template. The vector backbone was amplified with the primers CEBPb_bZIP_f (5′-CACC ACCACCATCACCTGTCCGATGAATACAAGATG-3′) and CEBPb_bZIP_r (5′-CGGTTTCTTTACCAGTCAGCAGTGGCCCGCCGAGG-3′). Both fragments were combined, digested with *Dpn*I and transformed into *E. coli* BL21(DE3). All constructs were sequenced before use and plasmids used for transfection experiments were purified with the EndoFree^®^ plasmid maxi kit (Qiagen, Hilden, Germany).

Plasmids were methylated using S*ss*I (New England Biolabs, Frankfurt am Main, Germany) as described previously (36). In brief, 10-20 μg plasmid DNA was incubated with S*ss*I (2.5 U/μg plasmid DNA) in the presence of 160 μM S-adenosylmethionine (SAM; New England Biolabs) for four hours at 37°C. Another 160 μM SAM was added after the first two hours of incubation. The unmethylated control DNA was treated as above without the addition of SAM. Following methylation DNA was purified and quantified spectrophotometrically. The completeness of methylation was controlled by digesting both methylated and unmethylated DNA with the methylation sensitive restriction enzyme *Hpy*CH4IV (New England Biolabs).

### Macrophage cell culture

The mouse macrophage cell line RAW264.7 (ATCC, TIB-71) was maintained in DMEM supplemented with 10% FCS, 1% glutamine, 100 units/ml penicillin, 100 μg/ml streptomycin (referred to as complete medium) at 37°C and 8% CO_2_. For isolation of primary bone marrow-derived macrophages (BMDMs), C57BL/6 mice were euthanized by CO_2_ inhalation. Femurs und tibias were excised and the bone marrow was flushed out with ice-cold PBS. Cells from multiple animals were pooled, suspended, and filtered through a cell strainer (Partec, Münster, Germany). BMDMs were differentiated by culturing bone marrow cells in complete medium and 20% L929 conditioned medium for eight days (fresh medium was added on day three) in bacteriological petri dishes or were frozen and stored at −80°C. Cells were harvested by being washed with cold PBS on day eight, seeded in appropriate cell culture dishes and used for experiments on day nine. Cells were stimulated with LPS as indicated in the respective figures. All animal experiments were approved by the appropriate ethical board (Niedersächsisches Landesamt für Verbraucherschutz und Lebensmittelsicherheit, Oldenburg, Germany).

### RNA isolation and absolute qRT-PCR

For absolute quantification of copy numbers by real-time PCR (qRT-PCR), total RNA was isolated using RNeasy®Mini Kit (Qiagen) and reverse described with M-MLV reverse transcriptase (Promega, Mannheim, Germany) and oligo-dT primer as described by the manufacturer. A ten-fold serial dilution series of the *Hind*III digested and purified pJet-Il36A plasmid, ranging from 6 × 10^4^ to 6 × 10^0^ copies/μl, was used to construct the standard curve. The concentration of the pJet-Il36A plasmid was measured photospectrometrically and the corresponding plasmid copy number was calculated as described previously (37).

RT-PCR was performed in a Stratagene™ Mx3005P qPCR instrument (Agilent Technologies, Waldbronn, Germany) using 10 μl QuantiTec SYBR Green PCR mix (Qiagen), 5 μl template DNA (either cDNA or plasmid DNA diluted as described above) and the primers Il36A_for and Il36A_rev (see plasmid construction) in a total volume of 20 μl. The PCR conditions were 95°C for 20 min, 40 cycles of 95°C for 20 s, 58°C for 30 s, and 72°C for 20 s, followed by a melting curve of the product as a control. Diluted standards were measured in triplicate and a standard curve was generated by plotting the threshold cycles (C*_t_*) against the natural log of the number of molecules. Based on the standard curve the number of *Il36A* cDNA molecules per 500 μg of total oligo-dT primed cDNA was calculated.

### Transient transfection and luciferase assay

Transient transfections were conducted using X-tremeGENE (Roche, Mannheim, Germany), according to the manufacturers’ protocol. RAW264.7 cells (1.0 × 10^5^/well) were co-transfected with the Firefly luciferase vector and control Renilla luciferase vector (pTK-RL) using 550 ng of DNA/reaction in total. Twenty-four hours after transfection, cells were treated with medium alone or 5.0 μg/ml LPS for additional eight hours. Cells were then lysed in 100 μl of 1× Passive Lysis Buffer (Promega) and luciferase activity in 10 μl aliquots of the cell lysates was measured using the Dual Luciferase Reporter Assay System (Promega) according to the manufacturer’s protocols. Firefly luciferase activity of individual cell lysates was normalized against Renilla luciferase activity.

### Electrophoretic mobility shift assays

For electrophoretic mobility shift assays (EMSA) equimolar amounts of complementary oligonucleotides were annealed and end labelled with [^32^P]dCTP using Klenow Fragment (3′→5′ exo-) (New England Biolabs). The following oligonucleotides were used: half-CRE•C/EBP_Il36A (5′-T CAGGTACTTCATCTTACGTCACCTAGT-3′, 5′-TCA GACTAGGTGACGTAAGATGAAGTAC-3′), methylated half-CRE•C/EBP_Il36A (5′-TCAGGTACTTCATCTTAm CGTCACCTAGT-3′, 5′-TCAGACTAGGTGAmCGTA AGATGAAGTAC-3′), consensus C/EBP (5′-TCAGCA GTCAGATTGCGCAATATCGGTC-3′, 5′-TCAGGAC CGATATTGCGCAATCTGACTG-3′). Nuclear extracts were prepared from RAW264.7 cells and primary mouse macrophages as described by Schreiber *et al*. (38) and band shift assays were performed exactly as described previously (5).

### DNA isolation and bisulphite sequencing

Genomic DNA was isolated using innuPREP Blood DNA Mini kit (Analytik Jena, Jena, Germany) according to the manufacturer’s instructions. The methylation profile of the proximal *Il36A* promoter was performed by bisulphite sequencing as described previously with minor modifications (39). In brief, 4 μg of genomic DNA was digested with *Bgl*II, purified by phenol-chloroform extraction and 500-1000 ng digested DNA was bisulphite-converted as described. Primers were designed using Methyl Primer Express®Software v1.0 (Applied Biosystems/ThermoFisher Scientific) to amplify specific regions of the genome following bisulphite conversion. The *Il36A* promoter region from –439 to –112 relative to the TSS on the (-)-strand was amplified using the primers Il36A_BSP_for (5′-GGAGGGTTTGTTAAGTATTTGT-3′) and Il36A_BSP_rev (5′-AATATCCACTAAAATCAACCTAAAA-3′). For bisulphite sequencing, PCR products were gel-purified and cloned into the pCR2.1-Topo Vector System (ThermoFisher Scientific) and sequenced using the M13rev primer (5′-CAGGAAACAGCTATGAC-3′). Sequencing results were analysed using QUMA software (40). Samples with conversion rate < 90% and sequences identity < 70% were excluded from the analysis. The minimum number of clones for each sequenced condition was ≥ 10.

### SDS-PAGE and Western blotting

For protein analysis by Western blotting, whole cell lysates were prepared in Cell Extraction Buffer (ThermoFisher Scientific) supplemented with protease inhibitor cocktail P8340, 1 mM AEBSF and Halt phosphatase inhibitor cocktail (ThermoFisher Scientific). Equal protein amounts (30 μg) were resolved on 12% BisTris-SDS-PAGE gels which were then blotted onto PVDF membranes. Membranes were blocked in 5% (w/v) skimmed milk for 1 h at room temperature, washed and incubated overnight with the respective primary antibodies at 4°C. After washing, the blots were incubated for 1 h with secondary antibodies against rabbit or goat IgG and washed. All washing steps were performed in TBS/0.05% (v/v) Tween® 20 (3 × 5 min). Blots were developed using SuperSignal West Pico Chemiluminescent Substrate (ThermoFisher Scientific) or AP-Juice Low Background (PJK GmbH, Kleinblittersdorf, Germany) and a ChemoCam Imager 3.2 (Intas, Göttingen, Germany). Densitometry was performed using LabImage 1D (Kapelan Bio-Imaging, Leipzig, Germany).

### Protein expression and purification

The DNA binding domain of mouse C/EBPβ was expressed as N-terminal 6×His fusion protein via pET45b in *E. coli* BL21(DE3). Bacteria were cultured in LB medium at 37°C to an optical density of A_600_ = 0.5. Protein production was induced by adding 1 mM isopropyl β-D-thiogalactopyranoside and the cultures were incubated at 30°C for 5 hours. Harvested bacteria were lysed using a French press (3 runs at 20.000 psi) in 1× LEW buffer (Macherey-Nagel, Düren, Germany) containing 5% (v/v) glycerol, 0.5 mM tris (2-carboxyethyl) phosphine (TCEP) and P8849 Protease Inhibitor Cocktail for His-tagged proteins (Sigma). The lysate was cleared at 25.000 × g for 30 min at 4°C and the supernatant was loaded on Protino®Ni-TED 2000 packed columns and purified as described by the manufacturer (Macherey-Nagel, Düren, Germany). The eluted protein was dialyzed against storage buffer (20 mM Tris-HCl pH 7.5, 150 mM NaCl, 5% (v/v) glycerol, 0.5 mM TCEP) and concentrated to approximately 5 ml using an Amicon® Ultra Centrifugal Filter Ultracel®-3K (UFC900324, Merck/Millipore, Darmstadt, Germany) before loading onto a HiTrap™-Heparin HP column (Cytiva, Marlborough, MA, United States). The Heparin column was then eluted using a step gradient of NaCl (250 mM to 2 M in 20 mM Tris-HCl pH 7.5, 5% (v/v) glycerol, 0.5 mM TCEP). The eluted fractions containing the purified protein were pooled and dialyzed against the storage buffer and concentrated as described above. The final concentration of the purified C/EBPβ-bZIP protein was estimated by Bradford protein assay (due to low number of aromatic residues).

### Fluorescence-based DNA binding assay

Fluorescence polarisation assays were performed using a GENios Pro microplate reader (Tecan, Männedorf, Switzerland) using the following oligonucleotides: FP_CREBmeth_f (5′-FAM-C ATCTTAmCGTCACCT-3′), FP_CREBmeth_r (5′AGGTGAmCGTAAGATG-3′), FP_CREB_f (5′-FAM-CATC TTACGTCACCT-3′), FP_CREB_r (5′AGGTGACGTA AGATG-3′). The annealed 6-carboxy-fluorescein (FAM)-labelled double-stranded oligonucleotides (5 nM) were incubated with an increasing amount of the protein for 30 min in 20 mM Tris-HCl pH 7.5, 150 mM NaCl, 5% (v/v) glycerol, 0.5 mM TCEP, 0.1 mg/ml BSA. Data were normalized and curves were fit individually using the neutcurve Python package (https://jbloomlab.github.io/neutcurve/). Averaged *K_D_* values and their standard deviations were reported.

### Data processing for visualization of *in vitro* C/EBPβ binding

We downloaded EpiSELEX-seq data for C/EBPβ (table containing the relative affinity of individual kmers from unmethylated and methylated libraries; GSE98652) (41) and visualized relative affinities of the kmers for the unmethylated library (Lib-U) and methylated library (Lib-M) with emphasis on half-CRE•C/EBP, consensus C/EBP and consensus CRE sequences. Energy logos were prepared by selecting all possible variations of the half-CRE•C/EBP sequence and subsequent visualization using the LogoGenerator of the REDUCE Suite v2.2 (http://reducesuite.bussemakerlab.org/index.php).

### Statistical analysis

Values are expressed as means ± SD. The exact sample size is given in the figure legends. Analysis using estimation statistics was done with Python 3.8 (Python Software Foundation, https://www.python.org/) and the DABEST package v0.3.1 (42). The generated Gardner-Altman estimation plots display the magnitude and robustness of the effect size and its bootstrapped 95% confidence interval (95% CI).

## Results

### Differential DNA methylation of a half-CRE•C/EBP element in the *Il36A* promoter in murine macrophages

We have previously shown that LPS-induced *Il36A* mRNA expression in murine macrophages is essentially mediated by binding of C/EBPβ to a half-CRE•C/EBP element within the *Il36A* promoter even though classical C/EBP recognition sites are present in the promoter sequence (5). In this study we aimed to dissect the specificity of C/EBPβ binding to the half-CRE•C/EBP element in relation to the level of methylation.

Detailed *in silico* analysis of the *Il36A* promoter region comprising 1120 bp upstream of the transcriptional start site revealed a GC content of 43.21%, the absence of any CpG islands but the presence of nine CpG dinucleotides (Fig. 1A). Isolation of genomic DNA and subsequent bisulfite sequencing of the region containing the CpG in the half-CRE•C/EBP element showed a low methylation of 9.70 ± 10.01% in RAW264.7 cells (Fig. 1B and D). In contrast, in the methylation level of this site in BMDMs was 66.95 ± 8.63% (Fig. 1C and D). The mean difference in methylation between both cell types is 57.2% (95% CI: 44.5, 67.5).

**Fig 1.**
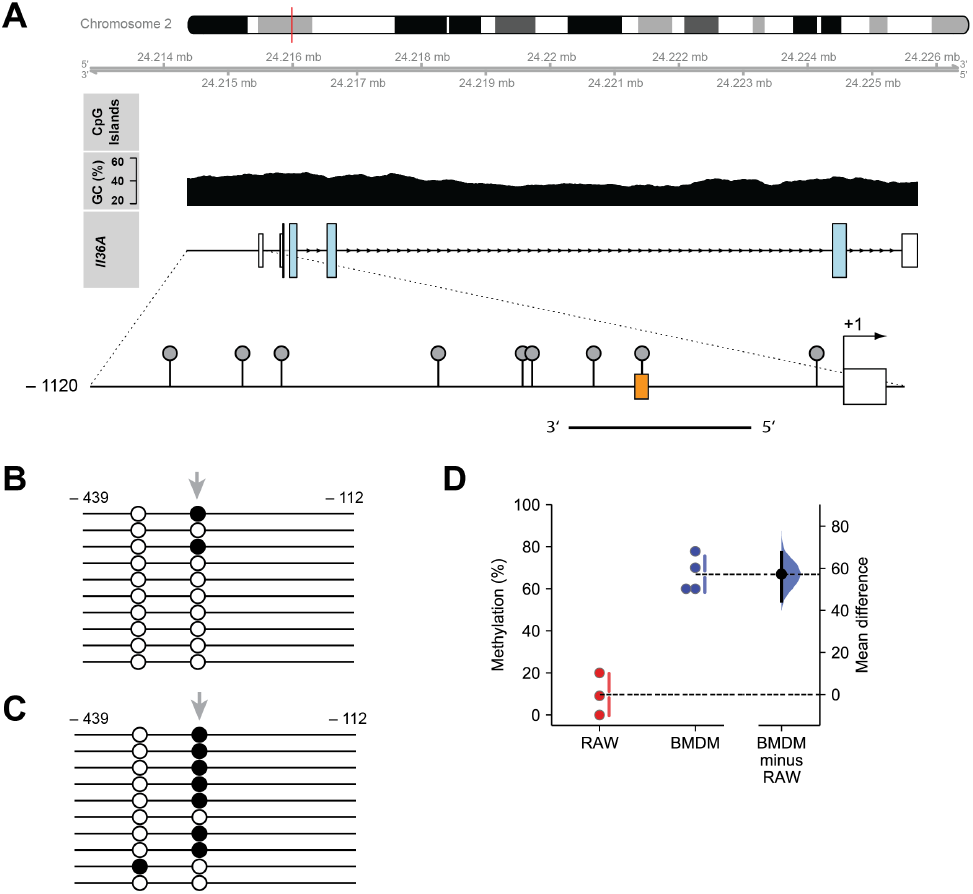
DNA methylation status of the half-CRE•C/EBP element in the *Il36A*promoter. **(A)** Schematic representation of the *Il36A* gene structure (Genebank entry KM205447.1), CpG islands, GC content (upper part) and the promoter region indicating CpG sites (lollipops, lower part). The position of *Il36A* on mouse chromosome 2 (red line), the half-CRE•C/EBP element (orange box) and the amplified region for bisulfite sequencing analysis on the (-) strand are indicated. **(B-C)** The DNA methylation status for the region indicated in (A) was determined in RAW264.7 **(B)** or BMDMs **(C)** by bisulfite sequencing analysis. Each line represents sequencing results of an individual clone (open circle, unmethylated CpG; filled circle, methylated CpG). The CpG in the half-CRE•C/EBP element is indicated by grey arrows. **(D)** Quantification of DNA methylation in RAW264.7 and BMDMs from n = 3 (RAW264.7) or n = 4 (BMDM) experiments. Individual data points and summary measurements (mean ± SD) are plotted on the left-hand side of the panel; effect size (mean differences, black dot) with bootstrapped 95% confidence intervals and resampling distribution are shown on the right-hand side of the panel.

These data show that the half-CRE•C/EBP motif in the *Il36A* promoter is differentially methylated in RAW264.7 cells and BMDMs, respectively.

### *In vitro* binding of C/EBPβ to the methylated half-CRE•C/EBP element in the *Il36A* promoter

Since it was demonstrated that CpG methylation of half-CRE elements enhances binding of C/EBP members and can activate tissue-specific gene expression the significant, differential methylation status of the CpG in the half-CRE•C/EBP might lead to differential binding activities of C/EBPβ in RAW264.7 macrophages and primary BMDMs. Therefore, we next analyzed binding of C/EBP and CREB members to the unmethylated and methylated half-CRE•C/EBP element in nuclear extracts by EMSA using a radiolabeled oligonucleotide spanning the region from −314 to −290 relative to the TSS of the murine *Il36A* gene.

We first studied the *in vitro* complex formation using nuclear extracts of RAW264.7 cells stimulated with LPS for 4 hours. In EMSA with the unmethylated probe, DNA-protein interactions appeared as two major complexes in unstimulated and LPS-stimulated cells. Whereas constitutive DNA binding activity of the slower migrating complexes was nearly unchanged (arrowheads, Fig. 2A), the binding activity of the faster migrating complexes increased in extracts from LPS-treated cells. Supershift experiments with antibodies against C/EBPβ, C/EBPδ, CREB-1, and ATF-4 indicated that only C/EBPβ and to a certain extent C/EBPδ are present in the inducible complex. Using the methylated probe, a similar inducible protein-DNA complex formation was observed. In comparison to the unmethylated probe, constitutive DNA binding activity was not detectable. The methylated probe also bound C/EBPβ and C/EBPδ but not CREB-1 nor ATF4 (Fig. 2A). Similar results were obtained with nuclear extracts from unstimulated and LPS-stimulated BMDMs. Irrespectively of the use of unmethylated or methylated probe LPS treatment induced a protein-DNA complex that supershifted with antibodies against C/EBPβ and to a minor extent by anti-C/EBPδ antibodies. Again, the formed complexes did not contain a detectable amount of CREB-1 nor ATF4 (Fig. 2B). However, in the extracts of both cell types the complexes formed with the methylated probe seemed to form more specific complexes with less background as compared to the unmethylated probe which is best emphasized by the reduced constitutive protein binding in the extracts from RAW264.7 macrophages using the methylated probe.

**Fig. 2.**
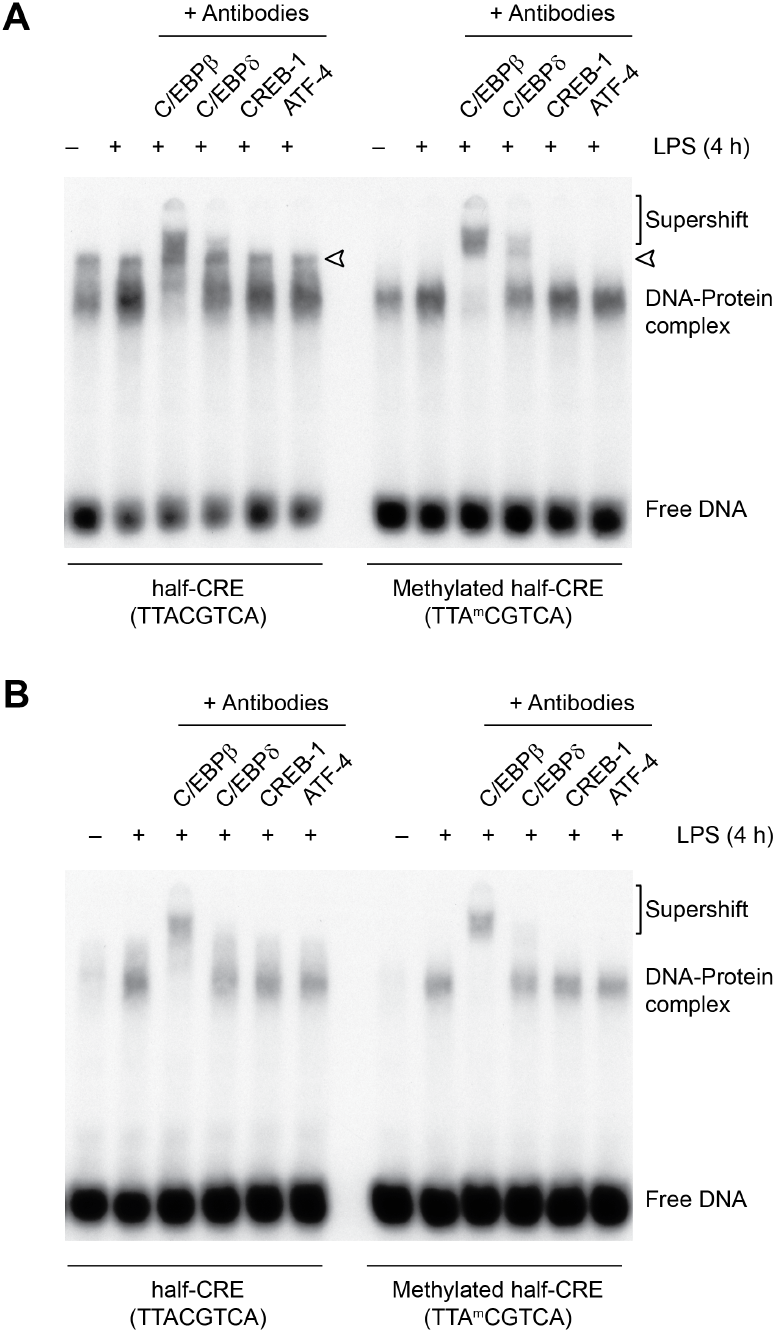
Transcription factor binding to the methylated half-CRE•C/EBP site of the *Il36A* promoter. **(A)** EMSAs were performed with nuclear extracts from untreated and LPS-treated (4 h, 5 μg/ml) RAW264.7 cells using radiolabeled oligonucleotides with unmethylated and methylated half-CRE•C/EBP element as probes. Supershift experiments were performed with specific antibodies against C/EBPβ, C/EBPδ, CREB-1/ATF-1, and ATF4. The arrowhead indicates a constitutively formed complex. One representative experiment of two independent experiments is shown. **(B)** EMSAs using the same probes and antibodies as in (A) performed with nuclear extracts from untreated and LPS-treated (4 h, 2.5 μg/ml) BMDMs. One representative experiment of two independent experiments is shown.

This suggested that C/EBPβ is able to bind unmethylated as well as methylated half-CRE•C/EBP sequences and that DNA methylation of the half-CRE•C/EBP element enhances specificity of protein binding.

### DNA binding of C/EBPβ is not enhanced by methylation

To test whether CpG methylation enhances binding of C/EBPs to DNA probes *in vitro* and to further analyze if there is any preferential binding to the unmethylated or methylated half-CRE•C/EBP oligonucleotides the protein-DNA complex formation was assessed with different approaches. First, we performed competition EMSA. For this, a radiolabeled oligonucleotide containing the C/EBP consensus motif (5′-TTGCGCAA-3′) was incubated with nuclear extracts of unstimulated and LPS-stimulated RAW264.7 cells in the presence of increasing amounts of unlabeled (cold) unmethylated or methylated half-CRE•C/EBP-Il36A oligonucleotides used before. As shown in Fig. 3A both oligonucleotides concentration-dependently decreased protein-DNA complex formation with comparable inhibitory effects.

**Fig. 3.**
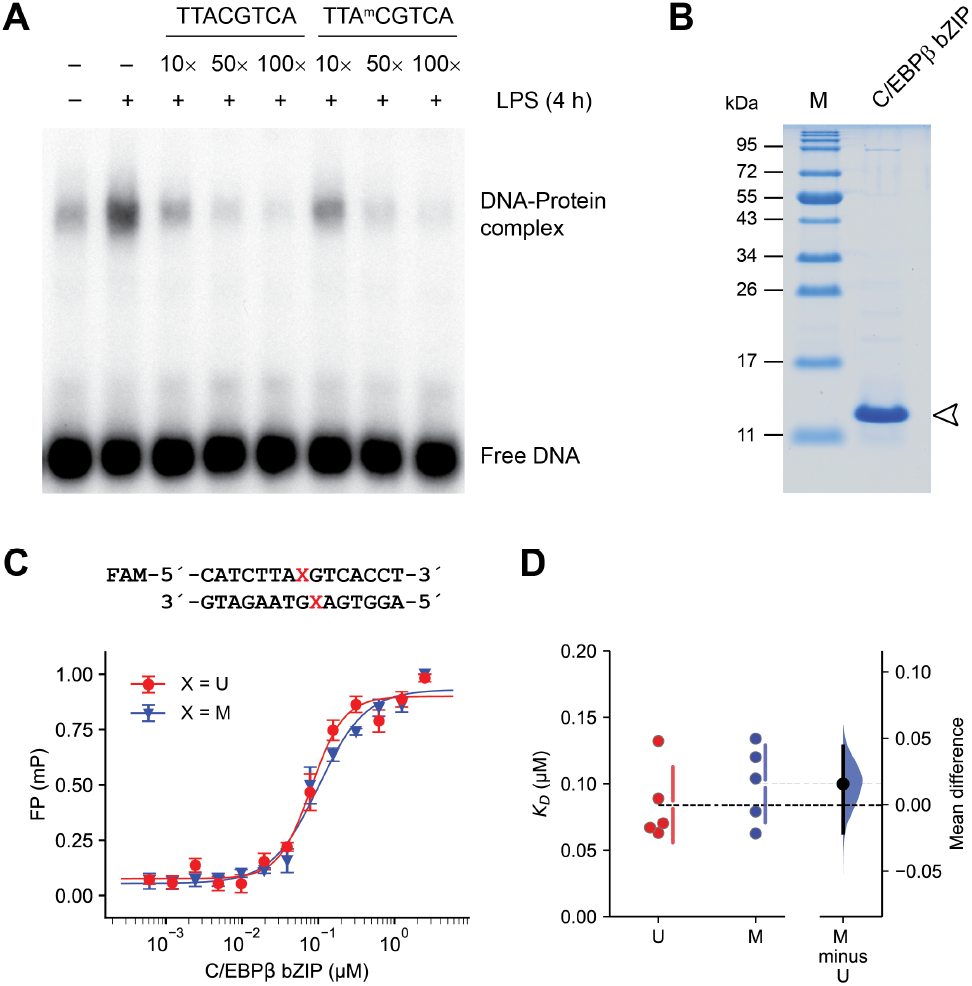
The effects of CpG methylation on DNA binding by C/EBPβ. **(A)** Competitive EMSAs were performed with a radiolabeled C/EBP consensus oligonucleotide probe and nuclear extracts from unstimulated and LPS-stimulated (4 h, 5 μg/ml) RAW264.7 cells. A 10-, 50-, and 100-fold molar excess of the unmethylated or methylated half-CRE•C/EBP-Il36A oligonucleotide was used in the competition reactions containing LPS-stimulated RAW264.7 nuclear extracts. One representative experiment of two independent experiments is shown. **(B)** 15% Bis-Tris PAGE gel showing the purified C/EBPβ-bZIP domain (2.5 μg, arrowhead) used in the fluorescence polarization (FP) assays. **(C)** DNA binding affinities of the C/EBPβ-bZIP domain to oligonucleotides of the *Il36A* half-CRE•C/EBP element containing a unmethylated (U) or methylated (M) central CpG measured by FP assays. Data represent mean ± SD from one representative experiments performed in triplicate. **(D)** Determination *K_D_* values using FP assays shown in (C) based on n = 5 experiments. Individual data points and summary measurements (mean ± SD) are plotted on the left-hand side of the panel; effect size (mean differences, black dot) with bootstrapped 95% confidence intervals and resampling distribution are shown on the right-hand side of the panel.

Next, we determined dissociation constants (*K_D_*) of the murine C/EBPβ-bZIP domain (Fig. 3B) to double-stranded half-CRE•C/EBP-Il36A oligonucleotides with unmethylated or methylated CpG using fluorescence polarization assays (Fig. 3C). The C/EBPβ DNA binding domain bound the unmodified oligonucleotides with a *K_D_* of 0.08 ± 0.03 μM (Fig. 3D). Under the same conditions, the methylated oligonucleotides were bound with a *K_D_* of 0.10 ± 0.03 μM (Fig. 3D). The mean difference between unmethylated and methylated oligonucleotides is 0.0157 μM (95% CI: −0.0213, 0.0438).

Together, these data further indicate, that C/EBPβ can recognize the half-CRE•C/EBP site located in the *Il36A* promoter irrespectively of the methylation status of central CpG *in vitro*.

### Methylation of the CpG in the half-CRE•C/EBP element does not inhibit *Il36A* promoter activity

Next, we assayed the relevance of CpG methylation of the half-CRE•C/EBP element in a system more resembling the *in vivo* situation in the nucleus. To evaluate whether the binding of C/EBPβ to the methylated half-CRE•C/EBP element is biologically relevant we performed transient transfections using RAW264.7 cells.

The *Il36A* promoter region, comprising the essential regulatory elements for gene induction between −357 and −45 relative to the TSS, was cloned into a reporter plasmid where all CpG dinucleotides have been deleted from the plasmid backbone (pCpGL) (36) (Fig. 4A). The use of a reporter without any CpGs in the parental plasmid backbone allowed evaluation of the effect of CpG methylation within the insert on transcriptional activation. The CpG in the half-CRE•C/EBP site in the pCpGL-*Il36A*^-357/-45^-Luciferase plasmid was enzymatically methylated *in vitro* using S*ss*I (CpG) methylase and S-adenosylmethionine. We confirmed methylation efficacy by restriction analysis. *Hpy*CH4IV linearized the unmethylated plasmid completely (asterisk, Fig. 4B) whereas the methylated plasmid was not linearized, indicating complete methylation (Fig. 4B).

**Fig. 4.**
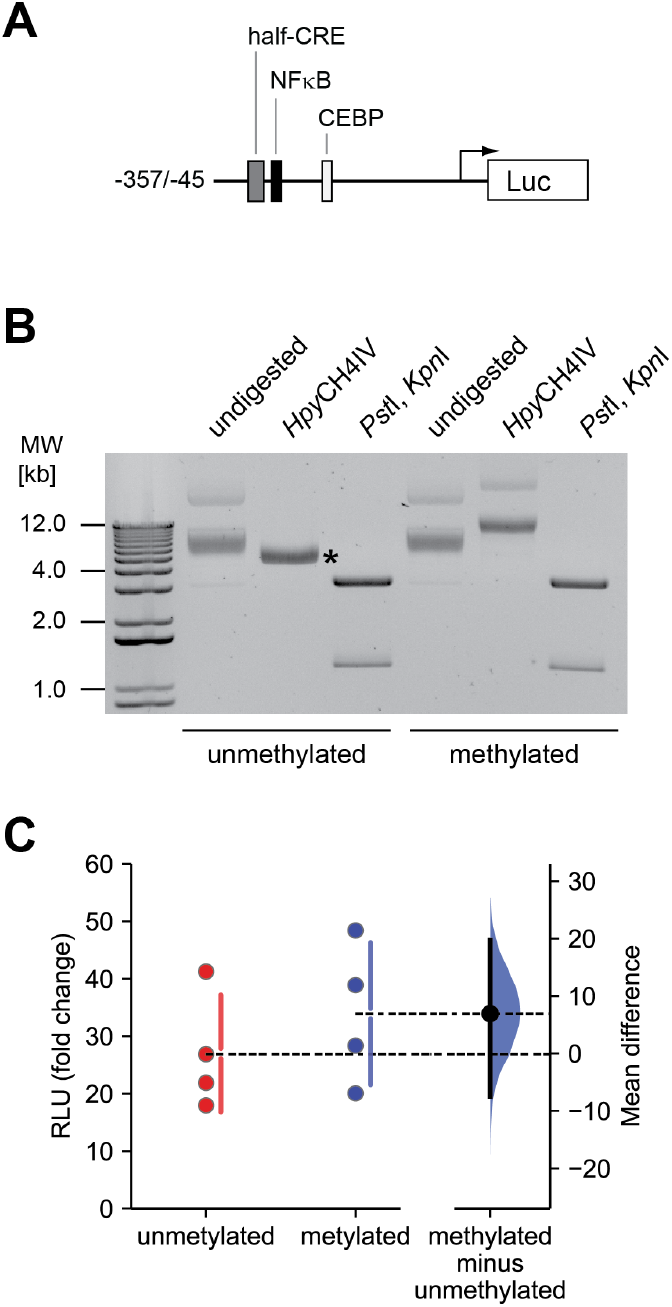
Inhibition of the *Il36A* promoter activity by methylation of the CpG sites in the half-CRE•C/EBP element. **(A)** Schematic representation of the pCpGL-*Il36A*^-357/-45^-Luciferase (Luc) reporter construct. Transcription factor binding sites are indicated. **(B)** *In vitro* methylation pCpGL-*Il36A*^-357/-45^ of using SssI CpG) methylase and S-adenosylmethionine. Methylation was confirmed by enzymatic digestion of the plasmid followed by gel electrophoresis. The unmethylated plasmid (lanes 1-3) is completely linearized by *Hpy*CH4IV (asterisk) whereas *Hpy*CH4IV does not linearize the methylated plasmid (lanes 4-6). Digestion with *Pst*I and *Kpn*I releases the *Il36A*^-357/-45^ insert from the plasmid **(C)** RAW 264.7 cells were co-transfected with the methylated/non-methylated pCpGL-*Il36A*^-357/-45^ construct along with the pRL-TK vector. 24 h after transfection cells were stimulated with LPS (100 ng/ml) for 8 h or left untreated. Firefly luciferase activity was normalized to that of Renilla luciferase and is expressed as fold change in relative luciferase induction (ratio LPS vs. ctrl) from n = 4 experiments. Individual data points and summary measurements (mean ± SD) are plotted on the left-hand side of the panel; effect size (mean differences, black dot) with bootstrapped 95% confidence intervals and resampling distribution are shown on the right-hand side of the panel.

Next, we transfected the unmethylated and methylated plasmids into RAW264.7 cells and determined luciferase activity in untreated cells or cells stimulated with 100 ng/ml LPS after 8 hours. As shown in Fig. 4C, stimulation with LPS led to an increased luciferase induction (fold change LPS vs. ctrl) in cells transfected with unmethylated and methylated reporter plasmid, respectively. The mean difference in luciferase induction between unmethylated and methylated plasmid is 6.94 (95% CI: −2027.47, 19.7).

These data show that CpG methylation of the half-CRE•C/EBP site does not significantly influence *Il36A* promoter activity in LPS-stimulated RAW264.7 cells.

### Methylation of the CpG in the half-CRE•C/EBP element does not influence Il-36α expression

Next, we analyzed whether the marginal expression differences in cells transfected with the methylated reporter plasmid were reflected by *Il36A* promoter activation in its ‘*in vivo*’ context in the nucleus.

For this we directly compared expression of *Il36A* induced by LPS in RAW264.7 cells and BMDMs, which have significantly different methylation levels of the half-CRE•C/EBP site (Fig. 1). Cells were stimulated with LPS for 8 hours and isolated total RNA was subjected to absolute quantification by qRT-PCR. As shown in Fig. 5A, *Il36A* induction was detectable after stimulation of RAW264.7 cells as well as BMDMs with LPS. We observed a mean difference in the fold change of *Il36A* mRNA induction of −101.4 (95% CI: −2276.3, 1748.2) between the two cell types.

**Fig. 5.**
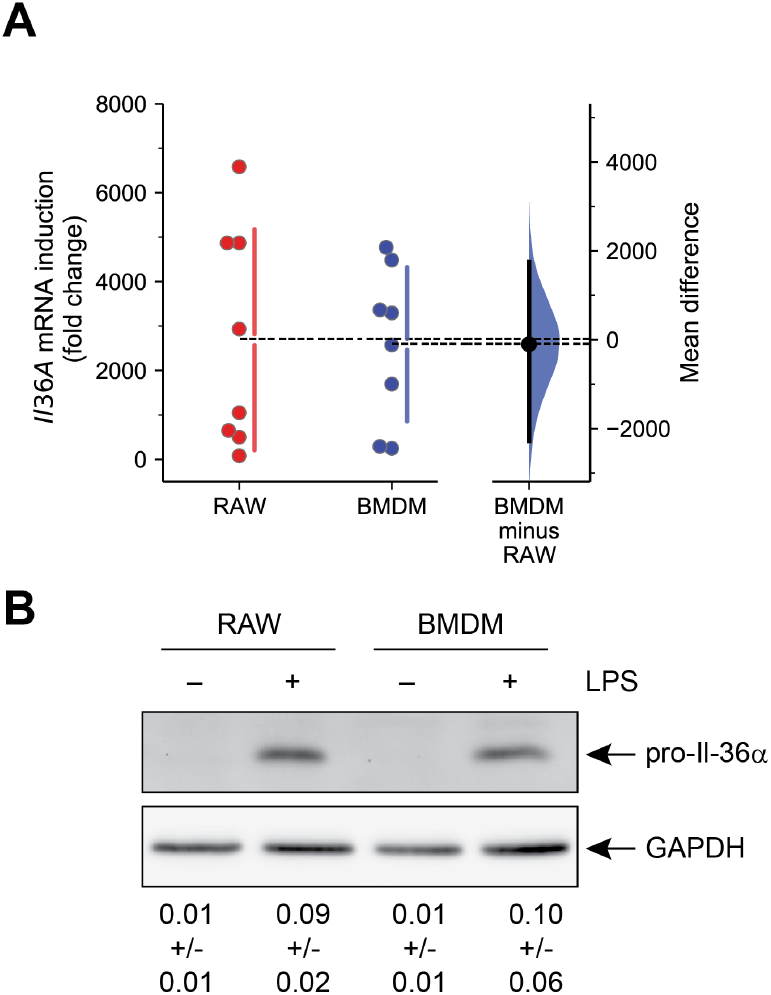
Induction of *Il36A* expression in LPS-stimulated macrophages. **(A)** RAW264.7 cells and BMDMs were stimulated with 1 μg/ml LPS for 8 h or left untreated and *Il36A* mRNA copy numbers were determined by qRT-PCR from n = 8 experiments and are expressed fold change in *Il36A* induction (ratio LPS vs. ctrl). Individual data points and summary measurements (mean ± SD) are plotted on the left-hand side of the panel; effect size (mean differences, black dot) with bootstrapped 95% confidence intervals and resampling distribution are shown on the right-hand side of the panel. **(B)** Western Blot analysis of pro-Il-36α expression in RAW264.7 cells and BMDMs stimulated with 1 μg/ml LPS for 8 h or left untreated. One representative Western blot out of 3 independent experiments is shown. Ratio of pro-Il-36α to GAPDH determined by densitometry is shown below (arbitrary units, n = 3 experiments, mean ± SD).

Furthermore, Western blot analysis of pro-Il-36α in cell lysates of RAW264.7 cells and BMDMs also revealed similar levels of the cytokine after stimulation with LPS (Fig. 5B) supporting the hypothesis that DNA methylation of the half-CRE•C/EBP site does not influence *Il36A* expression.

## Discussion

*Il36A* is expressed in a variety of cell types and in different tissues at different levels (11). Yet information on epigenetic regulation, in particular on CpG methylation and its possible impact on transcription factor binding are missing. We have previously shown that in murine macrophages LPS-induced *Il36A* mRNA expression requires binding of C/EBPβ to a half-CRE•C/EBP element within the *Il36A* promoter (5). Here we observed differential methylation of this element in RAW264.7 cells and primary murine macrophages, leading to the question whether this difference in methylation points to epigenetic regulation of *Il36A* by affecting TF binding and subsequent *Il36A* gene expression.

Regulation of gene expression by methylation of CpG motifs is one of several universal epigenetic mechanisms (43). The majority of data linking CpG methylation of the promoter regions and gene expression is derived from studies of genes with CpG islands (a region longer than 500 bp with an observed CpG/expected CpG ratio of 0.65) in the promoter region (44). CpG islands are not present in the *Il36A* promoter. However, the promoter harbours a number of CpGs proximal to the transcriptional start site, including one located within the essential half-CRE•C/EBP element. Such proximal, non-island CpGs are nowadays considered important for regulation of gene expression and examples of promoters affected by methylation of these non-island CpGs include e.g., *Il2, NOS2, MMP13, Il1B*, and *Il18BP* (45–48). While in most cases, methylation of the CpG sites results in transcriptional inactivation, for other genes methylation increased binding of TFs and thus resulted in stronger gene expression. Examples include *Bglap-rs1* and *MMP9* (24) as well as *PAX2* (49). This regulation at the epigenetic level may serve and control cell/tissue type-specific physiological functions or lead to reactivation of early developmental genes in malignancy.

Several lines of evidence presented herein suggest that methylation of the half-CRE•C/EBP element in the *Il36A* promoter has no influence on *Il36A* mRNA expression. Gel shift experiments and fluorescence polarization assays demonstrated that C/EBPβ is able to bind the unmethylated as well as methylated form of the CRE•C/EBP element *in vitro* which is agreement with earlier studies demonstrating binding of C/EBPβ to methylated consensus sequences (24). Interestingly, we also observed binding of C/EBPδ to both probes although overall binding was lower compared to C/EBPβ. This is in contrast to studies using methylation binding arrays showing that methylation inhibits binding of C/EBPδ (25). Furthermore, using an anti-ATF4 antibody in the supershift assays, we did not detect the formation of C/EBPβ/ATF4 heterodimers, indicating that the heterodimers are not formed after LPS stimulation because ATF4 is presumably not induced in sufficient amounts. However, for lysates from both cell types we detected less unspecific binding when the methylated probe was used, indicating that methylation interferes with binding of some unidentified TFs recognizing the unmethylated probe. Therefore, one could speculate that methylation modulates specificity of *Il36A* mRNA expression under certain circumstances. Indeed, Yin *et al*. demonstrated reduced binding for the majority of bZIP family members to their motifs upon methylation (Fig 5C in (16)). This supports the hypothesis that methylation of the half-CRE•C/EBP element in BMDMs may increase the specificity for C/EBPβ, one of the TFs in this family that tolerates methylation within the central CpG.

In contrast to the study by Rishi *et al*., we did not observe enhanced binding of C/EBPβ to the methylated motif *in vitro* (24). Since *Il36A* activation is C/EBPβ-dependent our data suggest that tolerance of a transcriptional regulator for CpG methylation helps to overcome otherwise functional epigenetic modifications. In line with this our findings are consistent with previous reports showing that *in vivo* binding by C/EBPβ tolerates CpG methylation (33). Furthermore, it is also in accordance with recently published data that compared relative affinities for C/EBPβ *in vitro* binding using methylated and unmethylated oligonucleotide libraries (41). Accordingly, we analyzed the specific sequence context of the *Il36A* half-CRE•C/EBP element by comparing methylated and unmethylated relative affinities with those of a consensus C/EBP motif and a consensus CRE motif, respectively based on published data for binding of C/EBPβ homodimers (41). Such a comparison revealed a relatively low binding affinity (compared to a consensus C/EBP motif) for the half-CRE•C/EBP motif that, however, did not differ between methylated and unmethylated sequences (Supplementary Fig. 1A). In contrast, *in vitro* binding of C/EBPβ to a consensus CRE motif was enhanced when methylated, confirming previous data (24). This methylation insensitivity is also reflected in the energy logos that are derived from the oligomer enrichment tables (Supplementary Fig 1B). In agreement, induction of luciferase expression upon LPS stimulation, the *Il36A* mRNA expression and the amount of pro-Il-36α at the protein levels did not differ significantly in the two cell types despite the observed difference in methylation. However, since only 66% of the cells in the BMDM population had a methylated half-CRE•C/EBP element, we cannot exclude that incomplete methylation influenced effect size of the qRT-PCR and Western Blot assays. Taken together, our data suggest that methylation of this motif does not influence endogenous promoter activity. Therefore, we speculate that asymmetry of the half-CRE•C/EBP element and/or the nucleotides flanking the central CpG in the different binding motifs might determine whether there is no effect or enhanced binding if the CpG is methylated.

Altogether, the data presented herein emphasize the potential of C/EBPβ to recognize methylated as well as unmethylated binding sites. Structural analysis or *in silico* modelling need to be performed in the future to unequivocally elucidate the binding mechanism of C/EBP to the half-CRE•C/EBP element present in the *Il36A* promoter. Furthermore, our data indicate that *Il36A* is most likely not regulated epigenetically by CpG methylation of the half-CRE•C/EBP element in the proximal promoter region of the gene in murine macrophages.

## ACKNOWLEDGEMENTS

We thank Michael Rehli (University Hospital Regensburg, Germany) for generously providing the pCpGL plasmid and Nanthapon Ruangkiattikul for assistance with the isolation of bone marrow cells. We thank the Henriques Lab (https://henriqueslab.github.io/) for the bioRχiv L^A^T_E_X template.

## AUTHOR CONTRIBUTIONS

Conceptualization, A.N. and R.G.; investigation, A.N. and N.J.; formal analysis, A.N. and R.G.; writing—original draft preparation, A.N and R.G.; writing—review and editing, all authors; funding acquisition, R.G. All authors have read and agreed to the published version of the manuscript.

## Supplementary Figures

**Supplementary Figure 1.**
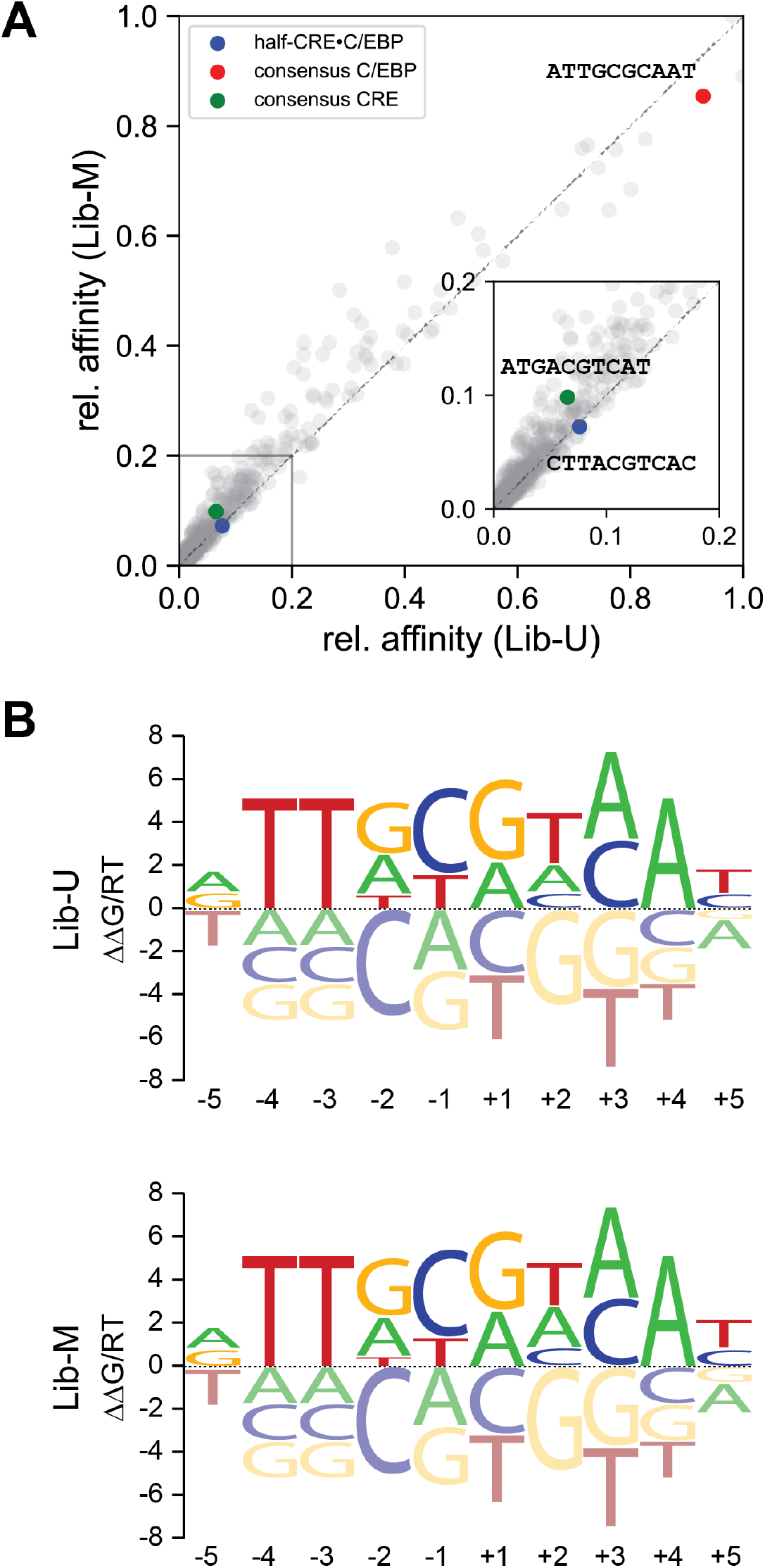
Methylation sensitivity of C/EBPβ based on GSE98652 data. **(A)** Comparison of the relative enrichment of 10-bp oligonucleotides between the unmethylated library (Lib-U) and the methylated library (Lib-M) for C/EBPb. The position of the half-CRE•C/EBP oligonucleotide is shown as blue dot, the position of the consensus C/EBP oligonucleotide is depicted in red and the position of the consensus CRE oligonucleotide is shown in green. **(B)** Energy logos for C/EBPb derived from Lib-U (upper logo) and Lib-M (lower logo).

## Bibliography

1. E.Y. Bassoy, J. E. Towne, and C. Gabay Regulation and function of interleukin-36 cytokines. Immunol Rev, 281(1):169–178, 2018. ISSN 0105-2896. doi: 10.1111/imr.12610.

2. S. Vigne, G. Palmer, C. Lamacchia, P. Martin, D. Talabot-Ayer, E. Rodriguez, F. Ronchi, F. Sallusto, H. Dinh, J. E. Sims, and C. Gabay. Il-36r ligands are potent regulators of dendritic and t cells. Blood, 118(22):5813–23, 2011. ISSN 0006-4971. doi: 10.1182/blood-2011-05-356873.

3. M. Bachmann, P. Scheiermann, L. Hardle, J. Pfeilschifter, and H. Muhl. Il-36gamma/il-1f9, an innate t-bet target in myeloid cells. J Biol Chem, 287(50):41684–96, 2012. ISSN 1083-351X (Electronic) 0021-9258 (Linking). doi: 10.1074/jbc.M112.385443.

4. A. M. Foster, J. Baliwag, C. S. Chen, A. M. Guzman, S. W. Stoll, J. E. Gudjonsson, N. L. Ward, and A. Johnston. Il-36 promotes myeloid cell infiltration, activation, and inflammatory activity in skin. J Immunol, 192(12):6053–61, 2014. ISSN 0022-1767. doi: 10.4049/jimmunol.1301481.

5. A. Nerlich, N. Ruangkiattikul, K. Laarmann, N. Janze, O. Dittrich-Breiholz, M. Kracht, and R. Goethe. C/ebpbeta is a transcriptional key regulator of il-36alpha in murine macrophages. Biochim Biophys Acta, 8:966–78, 2015. ISSN 0006-3002 (Print) 0006-3002 (Linking).

6. T. Aoyagi, M. W. Newstead, X. Zeng, S. L. Kunkel, M. Kaku, and T. J. Standiford. Il-36 receptor deletion attenuates lung injury and decreases mortality in murine influenza pneumonia. Mucosal Immunol, 10(4):1043–1055, 2017. ISSN 1933-0219. doi: 10.1038/mi.2016.107.

7. M. A. Kovach, B. Singer, G. Martinez-Colon, M. W. Newstead, X. Zeng, P. Mancuso, T. A. Moore, S. L. Kunkel, M. Peters-Golden, B. B. Moore, and T. J. Standiford. Il-36gamma is a crucial proximal component of protective type-1-mediated lung mucosal immunity in grampositive and -negative bacterial pneumonia. Mucosal Immunol, 10(5):1320–1334, 2017. ISSN 1933-0219. doi: 10.1038/mi.2016.130.

8. T. W. Lovenberg, P. D. Crowe, C. Liu, D. T. Chalmers, X. J. Liu, C. Liaw, W. Clevenger, T. Oltersdorf, E. B. De Souza, and R. A. Maki. Cloning of a cdna encoding a novel interleukin-1 receptor related protein (il 1r-rp2). J Neuroimmunol, 70(2):113–22, 1996. ISSN 0165-5728 (Print) 0165-5728.

9. S. Mutamba, A. Allison, Y. Mahida, P. Barrow, and N. Foster. Expression of il-1rrp2 by human myelomonocytic cells is unique to dcs and facilitates dc maturation by il-1f8 and il-1f9. Eur J Immunol, 42(3):607–17, 2012. ISSN 0014-2980. doi: 10.1002/eji.201142035.

10. R. Debets, J. C. Timans, B. Homey, S. Zurawski, T. R. Sana, S. Lo, J. Wagner, G. Edwards, T. Clifford, S. Menon, J. F. Bazan, and R. A. Kastelein. Two novel il-1 family members, il-1 delta and il-1 epsilon, function as an antagonist and agonist of nf-kappa b activation through the orphan il-1 receptor-related protein 2. J Immunol, 167(3):1440–6, 2001. ISSN 0022-1767 (Print) 0022-1767.

11. J. E. Towne, K. E. Garka, B. R. Renshaw, G. D. Virca, and J. E. Sims. Interleukin (il)-1f6, il-1f8, and il-1f9 signal through il-1rrp2 and il-1racp to activate the pathway leading to nf-kappab and mapks. J Biol Chem, 279(14):13677–88, 2004. ISSN 0021-9258 (Print) 0021-9258. doi: 10.1074/jbc.M400117200.

12. D. Schubeler. Function and information content of dna methylation. Nature, 517(7534): 321–6, 2015. ISSN 0028-0836. doi: 10.1038/nature14192.

13. F. Watt and P. L. Molloy. Cytosine methylation prevents binding to dna of a hela cell transcription factor required for optimal expression of the adenovirus major late promoter. Genes Dev, 2(9):1136–43, 1988. ISSN 0890-9369 (Print) 0890-9369.

14. S. M. Iguchi-Ariga and W. Schaffner. Cpg methylation of the camp-responsive enhancer/promoter sequence tgacgtca abolishes specific factor binding as well as transcriptional activation. Genes Dev, 3(5):612–9, 1989. ISSN 0890-9369 (Print) 0890-9369.

15. K. Gaston and M. Fried. Cpg methylation has differential effects on the binding of yy1 and ets proteins to the bi-directional promoter of the surf-1 and surf-2 genes. Nucleic Acids Res, 23(6):901–9, 1995. ISSN 0305-1048 (Print) 0305-1048.

16. Y. Yin, E. Morgunova, A. Jolma, E. Kaasinen, B. Sahu, S. Khund-Sayeed, P. K. Das, T. Kivioja, K. Dave, F. Zhong, K. R. Nitta, M. Taipale, A. Popov, P. A. Ginno, S. Domcke, J. Yan, D. Schubeler, C. Vinson, and J. Taipale. Impact of cytosine methylation on dna binding specificities of human transcription factors. Science, 356(6337), 2017. ISSN 0036-8075. doi: 10.1126/science.aaj2239.

17. X. Nan, H. H. Ng, C. A. Johnson, C. D. Laherty, B. M. Turner, R. N. Eisenman, and A. Bird. Transcriptional repression by the methyl-cpg-binding protein mecp2 involves a histone deacetylase complex. Nature, 393(6683):386–9, 1998. ISSN 0028-0836 (Print) 0028-0836. doi: 10.1038/30764.

18. Y. Liu, Y. O. Olanrewaju, Y. Zheng, H. Hashimoto, R. M. Blumenthal, X. Zhang, and X. Cheng. Structural basis for klf4 recognition of methylated dna. Nucleic Acids Res, 42(8): 4859–67, 2014. ISSN 0305-1048. doi: 10.1093/nar/gku134.

19. E. N. Nikolova, R. L. Stanfield, H. J. Dyson, and P. E. Wright. Ch...o hydrogen bonds mediate highly specific recognition of methylated cpg sites by the zinc finger protein kaiso. Biochemistry, 57(14):2109–2120, 2018. ISSN 0006-2960. doi: 10.1021/acs.biochem.8b00065.

20. T. Bartke, M. Vermeulen, B. Xhemalce, S. C. Robson, M. Mann, and T. Kouzarides. Nucleosome-interacting proteins regulated by dna and histone methylation. Cell, 143(3): 470–84, 2010. ISSN 0092-8674. doi: 10.1016/j.cell.2010.10.012.

21. C. G. Spruijt, F. Gnerlich, A. H. Smits, T Pfaffeneder, P. W. Jansen, C. Bauer, M. Munzel, M. Wagner, M. Muller, F Khan, H. C. Eberl, A. Mensinga, A. B. Brinkman, K. Lephikov, U. Muller, J. Walter, R. Boelens, H. van Ingen, H. Leonhardt, T. Carell, and M. Vermeulen. Dynamic readers for 5-(hydroxy)methylcytosine and its oxidized derivatives. Cell, 152(5): 1146–59, 2013. ISSN 0092-8674. doi: 10.1016/j.cell.2013.02.004.

22. G. C. Prendergast and E. B. Ziff. Methylation-sensitive sequence-specific dna binding by the c-myc basic region. Science, 251(4990):186–9, 1991. ISSN 0036-8075 (Print) 0036-8075.

23. S. Hu, J. Wan, Y. Su, Q. Song, Y. Zeng, H. N. Nguyen, J. Shin, E. Cox, H. S. Rho, C. Woodard, S. Xia, S. Liu, H. Lyu, G. L. Ming, H. Wade, H. Song, J. Qian, and H. Zhu. Dna methylation presents distinct binding sites for human transcription factors. Elife, 2: e00726, 2013. ISSN 2050-084x. doi: 10.7554/eLife.00726.

24. V. Rishi, P. Bhattacharya, R. Chatterjee, J. Rozenberg, J. Zhao, K. Glass, P. Fitzgerald, and C. Vinson. Cpg methylation of half-cre sequences creates c/ebpalpha binding sites that activate some tissue-specific genes. Proc Natl Acad Sci U S A, 107(47):20311–6, 2010. ISSN 1091-6490 (Electronic) 0027-8424 (Linking).

25. I. K. Mann, R. Chatterjee, J. Zhao, X. He, M. T. Weirauch, T. R. Hughes, and C. Vinson. Cg methylated microarrays identify a novel methylated sequence bound by the cebpb|atf4 heterodimer that is active in vivo. Genome Res, 23(6):988–97, 2013. ISSN 1549-5469 (Electronic) 1088-9051 (Linking).

26. J. Tsukada, Y. Yoshida, Y. Kominato, and P. E. Auron. The ccaat/enhancer (c/ebp) family of basic-leucine zipper (bzip) transcription factors is a multifaceted highly-regulated system for gene regulation. Cytokine, 54(1):6–19, 2011. ISSN 1096-0023 (Electronic) 1043-4666 (Linking).

27. T. Matsusaka, K. Fujikawa, Y. Nishio, N. Mukaida, K. Matsushima, T. Kishimoto, and S. Akira. Transcription factors nf-il6 and nf-kappa b synergistically activate transcription of the inflammatory cytokines, interleukin 6 and interleukin 8. Proc Natl Acad Sci U S A, 90 (21):10193–7, 1993. ISSN 0027-8424 (Print) 0027-8424.

28. S. E. Plevy, J. H. Gemberling, S. Hsu, A. J. Dorner, and S. T. Smale. Multiple control elements mediate activation of the murine and human interleukin 12 p40 promoters: evidence of functional synergy between c/ebp and rel proteins. Mol Cell Biol, 17(8):4572–88, 1997. ISSN 0270-7306 (Print) 0270-7306.

29. M. Matsumoto, Y. Sakao, and S. Akira. Inducible expression of nuclear factor il-6 increases endogenous gene expression of macrophage inflammatory protein-1 alpha, osteopontin and cd14 in a monocytic leukemia cell line. Int Immunol, 10(12):1825–35, 1998. ISSN 0953-8178 (Print) 0953-8178.

30. C. J. Lowenstein, E. W. Alley, P. Raval, A. M. Snowman, S. H. Snyder, S. W. Russell, and W. J. Murphy. Macrophage nitric oxide synthase gene: two upstream regions mediate induction by interferon gamma and lipopolysaccharide. Proc Natl Acad Sci U S A, 90(20): 9730–4, 1993. ISSN 0027-8424 (Print) 0027-8424.

31. D. J. Wadleigh, S. T. Reddy, E. Kopp, S. Ghosh, and H. R. Herschman. Transcriptional activation of the cyclooxygenase-2 gene in endotoxin-treated raw 264.7 macrophages. J Biol Chem, 275(9):6259–66, 2000. ISSN 0021-9258 (Print) 0021-9258.

32. J. R. Newman and A. E. Keating. Comprehensive identification of human bzip interactions with coiled-coil arrays. Science, 300(5628):2097–101, 2003. ISSN 0036-8075. doi: 10.1126/science.1084648.

33. H. Zhu, G. Wang, and J. Qian. Transcription factors as readers and effectors of dna methylation. Nat Rev Genet, 17(9):551–65, 2016. ISSN 1471-0056. doi: 10.1038/nrg.2016.83.

34. J. Yang, J. R. Horton, D. Wang, R. Ren, J. Li, D. Sun, Y. Huang, X. Zhang, R. M. Blumenthal, and X. Cheng. Structural basis for effects of cpa modifications on c/ebpbeta binding of dna. Nucleic Acids Res, 47(4):1774–1785, 2019. ISSN 1362-4962 (Electronic) 0305-1048 (Linking). doi: 10.1093/nar/gky1264.

35. J. García-Nafría, J. F. Watson, and I. H. Greger. Iva cloning: A single-tube universal cloning system exploiting bacterial in vivo assembly. Sci Rep, 6:27459, 2016. ISSN 2045-2322. doi: 10.1038/srep27459.

36. M. Klug and M. Rehli. Functional analysis of promoter cpg methylation using a cpg-free luciferase reporter vector. Epigenetics, 1(3):127–30, 2006. ISSN 1559-2294.

37. J. A. Whelan, N. B. Russell, and M. A. Whelan. A method for the absolute quantification of cdna using real-time pcr. J Immunol Methods, 278(1-2):261–9, 2003. ISSN 0022-1759 (Print) 0022-1759.

38. E. Schreiber, P. Matthias, M. M. Müller, and W. Schaffner. Rapid detection of octamer binding proteins with ‘mini-extracts’, prepared from a small number of cells. Nucleic Acids Res, 17(15):6419, 1989. ISSN 0305-1048 (Print) 0305-1048. doi: 10.1093/nar/17.15.6419.

39. S. J. Clark, A. Statham, C. Stirzaker, P. L. Molloy, and M. Frommer. Dna methylation: bisulphite modification and analysis. Nat Protoc, 1(5):2353–64, 2006. ISSN 1750-2799. doi: 10.1038/nprot.2006.324.

40. Y. Kumaki, M. Oda, and M. Okano. Quma: quantification tool for methylation analysis. Nucleic Acids Res, 36(Web Server issue):W170–5, 2008. ISSN 0305-1048. doi: 10.1093/nar/gkn294.

41. J. F. Kribelbauer, O. Laptenko, S. Chen, G. D. Martini, W. A. Freed-Pastor, C. Prives, R. S. Mann, and H. J. Bussemaker. Quantitative analysis of the dna methylation sensitivity of transcription factor complexes. Cell Rep, 19(11):2383–2395, 2017. doi: 10.1016/j.celrep.2017.05.069.

42. J. Ho, T. Tumkaya, S. Aryal, H. Choi, and A. Claridge-Chang. Moving beyond p values: data analysis with estimation graphics. Nat Methods, 16(7):565–566, 2019. ISSN 1548-7091. doi: 10.1038/s41592-019-0470-3.

43. C. D. Allis and T. Jenuwein. The molecular hallmarks of epigenetic control. Nat Rev Genet, 17(8):487–500, 2016. ISSN 1471-0056. doi: 10.1038/nrg.2016.59.

44. D. Takai and P. A. Jones. Comprehensive analysis of cpg islands in human chromosomes 21 and 22. Proc Natl Acad Sci U S A, 99(6):3740–5, 2002. ISSN 0027-8424 (Print) 0027-8424. doi: 10.1073/pnas.052410099.

45. J. K. Northrop, R. M. Thomas, A. D. Wells, and H. Shen. Epigenetic remodeling of the il-2 and ifn-gamma loci in memory cd8 t cells is influenced by cd4 t cells. J Immunol, 177(2): 1062–9, 2006. ISSN 0022-1767 (Print) 0022-1767. doi: 10.4049/jimmunol.177.2.1062.

46. K. Hashimoto, M. Otero, K. Imagawa, M. C. de Andrés, J. M. Coico, H. I. Roach, R. O. Oreffo, K. B. Marcu, and M. B. Goldring. Regulated transcription of human matrix metalloproteinase 13 (mmp13) and interleukin-1 β (il1b) genes in chondrocytes depends on methylation of specific proximal promoter cpg sites. J Biol Chem, 288(14):10061–72, 2013. ISSN 0021-9258 (Print) 0021-9258. doi: 10.1074/jbc.M112.421156.

47. T. J. Gross, K. Kremens, L. S. Powers, B. Brink, T. Knutson, F. E. Domann, R. A. Philibert, M. M. Milhem, and M. M. Monick. Epigenetic silencing of the human nos2 gene: rethinking the role of nitric oxide in human macrophage inflammatory responses. J Immunol, 192(5): 2326–38, 2014. ISSN 0022-1767. doi: 10.4049/jimmunol.1301758.

48. M. Bachmann, J. Pfeilschifter, and H. Muhl. Epigenetic regulation by cpg methylation splits strong from retarded ifngamma-induced il-18bp in epithelial versus monocytic cells. Biochim Biophys Acta Gene Regul Mech, 1861(3):191–199, 2018. ISSN 1874-9399 (Print) 1874-9399. doi: 10.1016/j.bbagrm.2018.01.020.

49. N. Jia, J. Wang, Q. Li, X. Tao, K. Chang, K. Hua, Y. Yu, K. K. Wong, and W. Feng. Dna methylation promotes paired box 2 expression via myeloid zinc finger 1 in endometrial cancer. Oncotarget, 7(51):84785–84797, 2016. ISSN 1949-2553. doi: 10.18632/oncotarget.12626.

